# Intraspecific variation in Artiodactyla social organisation: A Bayesian phylogenetic multilevel analysis of detailed population-level data

**DOI:** 10.1101/603399

**Authors:** M.I. Miles, A.V. Jaeggi, M. Festa-Bianchet, C. Schradin, L.D. Hayes

## Abstract

Understanding *inter*-specific variation in social systems is a major goal of behavioural ecology. Previous comparative studies of mammalian social organisation produced inconsistent results, possibly because they ignored *intra*-specific variation in social organisation (IVSO). The Artiodactyla have been the focus of many comparative studies as they occupy a wide diversity of habitats and exhibit large variation in life history patterns as well as other potential correlates of social organisation. Here we present the first systematic data on IVSO among Artiodactyla, infer their ancestral social organisation, and test whether habitat, sexual dimorphism, seasonal breeding, and body size predict inter- and intraspecific variation in social organisation. We found data on social organisation for 110 of 226 artiodactyl species, of which 74.5% showed IVSO. Using Bayesian phylogenetic multilevel models, the ancestral artiodactyl population was predicted to have a variable social organisation with significantly higher probability (0.77, 95% CI 0.29-1.00) than any non-variable form (i.e. solitary, pair-living, group-living). Greater sexual dimorphism and smaller body size both predicted more IVSO; smaller body size also predicted a higher likelihood of pair-living. Our results challenge the long-held assumption that ancestral Artiodactyla were pair-living and strongly imply that taking IVSO into account is crucial for understanding mammalian social evolution.

## 1. Introduction

Animals show remarkable *inter*-specific variation in social systems [1, 2], and understanding the sources of this diversity is a major goal of behavioural ecology. Social systems are characterized by four components [3, 4]: i) social organisation: the size, sex-age, and kin composition of groups, ii) mating system, iii) social structure: relationships emerging from repeated interactions among individuals, and iv) parental and allo-parental care. These components are interdependent. For example, the number and spatial distribution of individuals characterize their social organisation but also constrain their mating tactics [4].

There have been numerous comparative analyses of mammalian social organisation [6–8]. However, inconsistent results have emerged from these studies for several taxa, including primates and carnivorans. In primates, it has been suggested that pair-living species evolved exclusively from solitary [7] or from both solitary and group-living ancestors [8, 9]. In carnivorans, the long-held hypothesis that social evolution involved transitions in social organisation from a solitary ancestor into more advanced forms of group living [solitary ancestor hypothesis: 10, 11] has been questioned [6].

These inconsistent results likely occurred for several reasons. First, studies have relied on different datasets, methods of analysis, and conceptual frameworks [3]. In an effort to account for as many species as possible, some studies relied on information from secondary sources and taxonomic inference, such as the untested assumption that members of the same genus share the same social organisation [12]. Other studies used confusing terminology or failed to distinguish between social organisation and mating system [3]. For example, some studies inferred monogamy (mating system) from the observation of male-female pairs (social organisation) [9, 13]. To resolve these issues, comparative studies should rely exclusively on data from primary sources, and distinguish social organisation from other social system components.

Most comparative studies of mammalian social organisation relied on a single trait value for each species, yet social systems can be dynamic [14–16]. *Intra*specific variation in social organisation (IVSO) occurs when adults of a species show two or more forms of social organisation [6, 14, 17]. Variation can also occur in the composition of groups, such that a species may live in different types of groups, e.g., unisex vs. multi-sex groups. In mammals, IVSO has been reported in numerous species from different orders [6, 18–22], transforming our understanding of mammalian social evolution. For example, in carnivorans (Order: Carnivora) and shrews (Order: Euliptophya), it was long believed that the ancestral state was solitary. However, a variable ancestral state was found to be equally likely after taking IVSO into account (Carnivora: [6]; Euliptophya: [21]). More broadly, ignoring intraspecific variation can increase statistical type II error rates [23–25] and lead to spurious conclusions about social evolution [17, 26]. Modern comparative methods such as phylogenetic mixed-effects (a.k.a. multilevel) models or measurement-error models [24, 27] can easily incorporate intraspecific variation. Thus, comparative studies should include intraspecific variation in social organisation.

IVSO may arise, for example, if individuals of both sexes can respond to unpredictable or changing ecological conditions by changing their social tactic [16, 17]. Further, IVSO may vary with body size, which correlates strongly with life-history pace [34] as well as available anti-predator strategies [35]. Intraspecific variation in group size and composition is also expected in seasonal breeders. During the breeding season, reproductive competition can exclude some individuals from groups, thus causing IVSO [36]. Alternatively, relaxed competition during the non-breeding season may allow the formation of larger groups, particularly if grouping has survival benefits [e.g., anti-predator strategies; 18, 37]. Thus, we expect greater variability in social organisation among species occupying a wider range of habitats and among seasonal breeders.

The order Artiodactyla is well suited for comparative studies of social evolution because its members exhibit both inter- and intraspecific variation in social organisation, occupy a wide diversity of habitat types, and exhibit a range of body sizes, sexual dimorphism, as well as both seasonal and non-seasonal breeding [18, 38]. Habitat heterogeneity and availability of protective cover are associated with *inter*specific variation in social organisation of many artiodactyls [18]. Generally, groups are larger in open areas [18, 39], with solitary species mostly living in dense forests [40]. Group-living and large body size are adaptations to open habitats characterized by high predation risk [41]. Sexual dimorphism in body size and seasonal breeding, common in artiodactyls, are also associated with interspecific variation in social organisation [18]. Most sexually dimorphic species live in unisex groups or as solitary individuals, forming mixed sex groups only in the breeding season [42, 43], [but see 44, 45]. Monomorphic species live alone, in pairs, or mixed sex groups [46]. Sexual dimorphism is also correlated with body size [18]. We therefore expected these factors to influence IVSO.

We first describe *inter*specific and *intra*specific variation in artiodactyl social organisation, using only data from published studies on wild populations. Our second objective was to infer the ancestral social organisation of artiodactyls. We used a detailed phylogeny and modern comparative methods to evaluate competing hypotheses about artiodactyl social evolution, namely 1) from pair-living to group-living [7] or 2) from IVSO to single types of social organisation [6]. Our third objective was to determine the extent to which habitat, sexual dimorphism, body size and breeding seasonality predict variation in social organisation. Specifically, we predicted that the likelihood of IVSO and the total number of social organisations in a species (i) increase with greater number of habitats, (ii) decrease in open habitats due to predation pressure favoring group-living, (iii) sexually dimorphic than monomorphic species and (iv) are greater in seasonal breeders than non-seasonal breeders.

## 2. Methods

### (a) Data collection

Searches were conducted using Web of Science and Google Scholar to find primary sources reporting social organisation for all 226 extant species of Artiodactyla [47]. The initial search consisted of the scientific name (genus and species) and a keyword (‘social’, ‘herd’, or ‘group’). If no sources were found, a final search used only the scientific name. In *Web of Science*, search results were refined by selecting three research areas: ‘zoology’, ‘behavioral science’, and ‘environmental science/ecology’, and document type ‘article’. Lab-based studies, studies in enclosures smaller than 1,000-hectares, and studies that included manipulation of individuals, groups, or resources were discarded. From the 267 primary sources, we coded the following social organisation(s): 1) Solitary, 2) Pair-living, 3) Sex-specific social organisation 4) Unisex groups, 5) Single female/multi-male, 6) Multi-female/single male, and 7) Multi-female/multi-male (Supplementary Material Table S1).

### (b) Determining variable social organisation

Variable social organisation was identified when 1) both sexes had more than one form of social organisation in the same population [e.g., solitary and pair-living; 17] or between populations or 2) multiple types of groups occurred within the same population or in different populations (e.g., FFM and FFMM groups). For statistical analyses, overall population-level social organisations were categorized as: 1) Solitary, 2) Pair-living, 3) Sex-specific, 4) Non-variable group-living, and 5) variable, including populations with multiple forms of group-living. We categorized cases where one sex was solitary and the other was in unisex groups as a specific form of social organisation (sex-specific) and cases in which both sexes lived in unisex groups as a form of group-living. Additional details are provided in Supplementary Material S2.

### (c) Predictor variables

Each species was categorized as either seasonal or non-seasonal breeder [38]. The extent of sexual dimorphism was calculated as the ratio of adult male to female body mass using data reported in Pérez-Barbería & Gordon [38]. Body size was included as mean adult female body mass. Habitat type was derived from the primary source and categorized based on IUCN classification (www.iucn.org) as desert, forest, rocky areas, savanna, grassland, shrubland, wetlands, or artificial.

### (d) Phylogeny

We used the mammal supertree from Bininda-Emonds et al. [48]. Some species names in the database had to be amended to match the phylogeny as detailed in the accompanying R code. In virtually all cases, a name mismatch could be resolved by finding a pseudonym for that species through www.iucn.org, or by using a sister species that was not included in the database. In one case, two closely-related taxa missing from the supertree (*Moschus leucogaster* and *Moschus cupreus*) were proxied by the same sister species (*Moschus chrysogaster*).

### (e) Statistical analysis

We used Bayesian phylogenetic mixed-effects models, accounting for the multilevel structure of the data (populations nested within species) and the phylogenetic relationships among species [24, 27]. Predictors included sexual dimorphism, female body size, breeding seasonality, and number of habitats. Type of habitat was modeled as a random intercept. All models controlled for research effort by including the number of studies. To control for potential geographical biases continent was included as a random intercept.

Prior to fitting the model, we estimated the likely ancestral state for body size, sexual dimorphism, and breeding seasonality (see Supplementary Material 3). We then centered body size and sexual dimorphism on these estimated ancestral states and chose the likely ancestral breeding seasonality as the reference category. Consequently, the estimated ancestral social organisation (i.e. the global intercepts of the multilevel models) is contingent on predictors that are *also* at their likely ancestral state.

To model the likelihood of several mutually exclusive categorical traits (i.e. different social organisations) and how the likelihood of each trait was affected by other variables we used a multinomial model [52] (see Supplementary Material 3). We highlight any covariates that influence the likelihood of different social organisations, and thus may explain evolutionary transitions from the ancestral state. Additional details are in Supplementary Material 4.

We fit all models in a Bayesian framework [53] in Stan [54] through the RStan interface [55] using *brms* v. 2.5.1. [56]. Bayesian estimation produces a posterior probability distribution for each parameter, which can be summarized in various ways; here we report the mean and 95% credible intervals and occasionally the proportion of the posterior that lies above or below a certain value (“PP”). Phylogenetic signal was calculated as the proportion of variance captured by the phylogenetic random effect(s) following [57]. All models converged as assessed with the potential scale reduction factor (all =<1.01), effective sample sizes (all >500), and by visually examining trace plots of the Markov chains. Details on model fitting can be gleaned from the accompanying R code (available at https://github.com/adrianjaeggi/artiodactyl.socialorg).

## 3. Results

We found data on social organisation for 247 populations from 110 of the 226 extant artiodactyl species. The majority of these species showed variation in their social organisation at the species level (74.5%, 82 out of 110). Five species were strictly solitary (4.5%), only one was strictly pair-living (0.9%), one showed sex-specific social organisation (0.9%), and eleven showed only one form of group-living (Table 1). A more detailed breakdown of variable social organisation is available in Supplementary Material Table S5. At the population level, 62% (155/247) of all populations also had variable social organisation. Of the 82 species showing variable social organisation, five (6.1%) showed variation between populations, twenty-nine (35.4%) showed variation within a population, and forty-eight (58.5%) showed variable social organisation both between populations and within a population.

**Table 1.** Social organisations of Artiodactyla species

| **Category** | **No. and percentage** |
| --- | --- |
| **Solitary** | 5 (4.5%) |
| **Pair-living** | 1 (0.9%) |
| **Sex-specific** | 1 (0.9%) |
| **Group-living, only, but split into:** |  |
| - No variation in group composition | 11 (10.0%) |
| - Variable group composition* | 21 (19.1%) |
| - Unknown composition | 10 (9.1%) |
| **Species with more than one social organisation** | 61 (55.5%) |
| **Species with variable group composition** | 42 (38.2%) |
| **Species with any form of IVSO, including variable group composition** | 82 (74.5%) |
| **Species with data** | 110 (48.7%) |
| **Species with no data** | 116 (51.3%) |

\* Variable includes more than one of the following: single male, multiple females (MFF); single female, multiple males (FFM); multiple males, multiple females (MMFF); unisex groups, both male-only and female-only in the same population.

A summary of the phylogenetic multilevel model is available in the Supplementary Material S6. The intercepts reflect a non-seasonally-breeding species of ancestral body size and sexual dimorphism that lives in only one habitat and was studied once. An ancestral population with these characteristics was predicted to have a variable social organisation with significantly higher probability (0.77, 95% CI 0.29-1.00) than any non-variable form (Figure 1).

**Figure 1.**
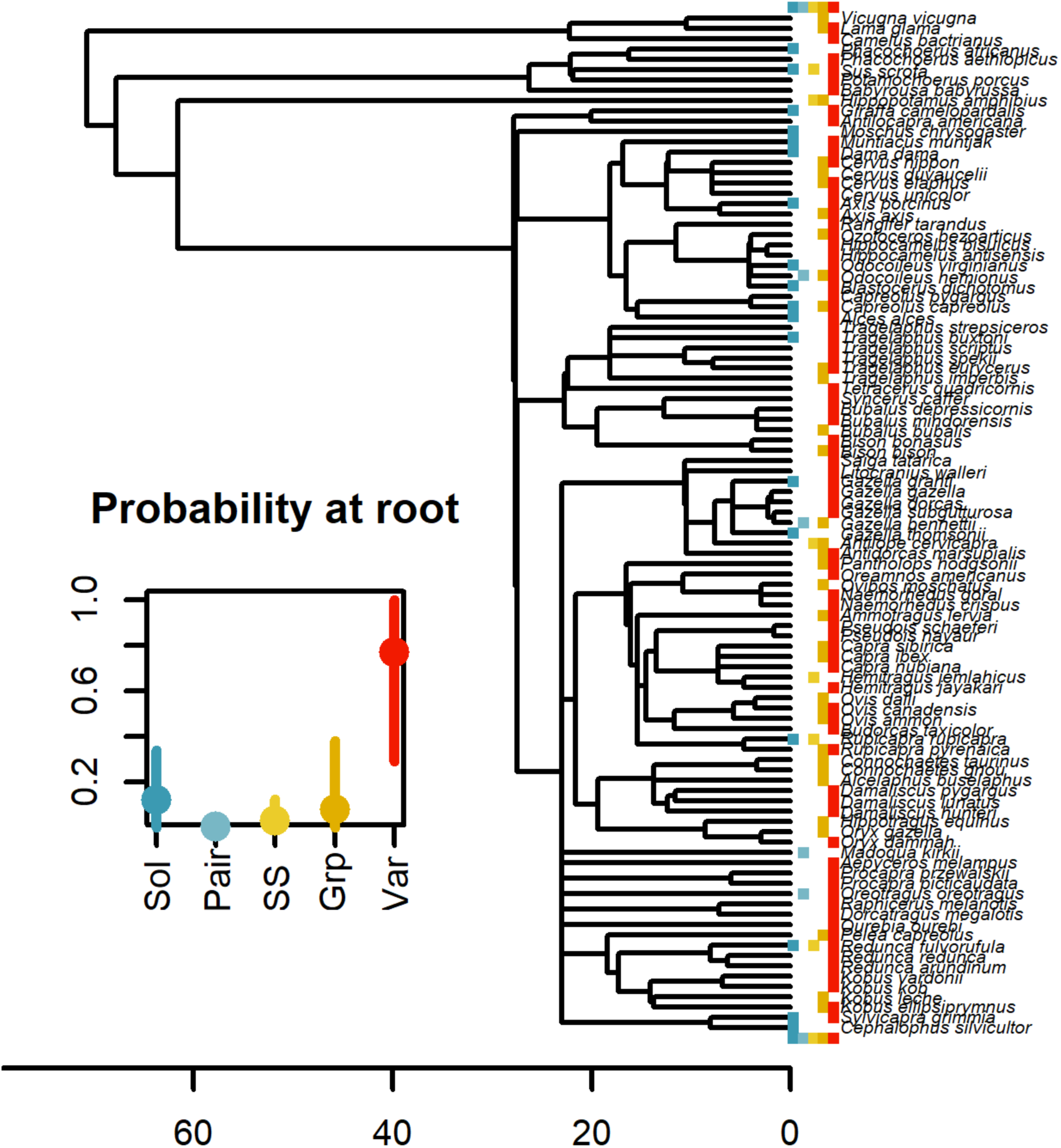
Phylogeny of artiodactyls with data on social organisation, along with the probability of each social organisation for the last common ancestor (Sol=Solitary, Pair=Pair-living, SS=Sex-specific, Grp=Non-variable group-living, Var=Variable). The colored boxes at the tips of the phylogeny show social organisations observed in populations of extant species. The five possible states (solitary, pair-living, sex-specific, group-living, variable) are plotted above and below the phylogeny in this order and the same colors as the inserted figure on ancestral social organisation. The scale bar shows million years before present.

The likelihood of variable social organisation increased with degree of sexual dimorphism (odds ratio for 1SD change = 2.91, 95% CI = 1.16 – 8.94), and decreased as female body mass increased (odds ratio for 1SD change = 0.39, 95% CI = 0.16 – 0.80; Figure 2). Pair-living was more likely with lower female body mass (Figure 2). Unsurprisingly, the probability of variable and sex-specific social organisations increased with study effort. No other associations were “significant” at the 95% CI level, but transitions to group-living were likelier with greater sexual dimorphism (PP=0.85) and seasonal breeding (PP=0.87). In terms of habitat type, the prediction of variable social organisation being less likely in open (savanna and native grasslands) than closed (forest) habitats was not supported (PP=0.37). Similarly, support for group-living being more likely in open habitats was weak (PP=0.63). The phylogenetic signal in social organisation was weak but greater than 0 (mean = 0.24, 95% CI = 0.05 – 0.48).

**Figure 2.**
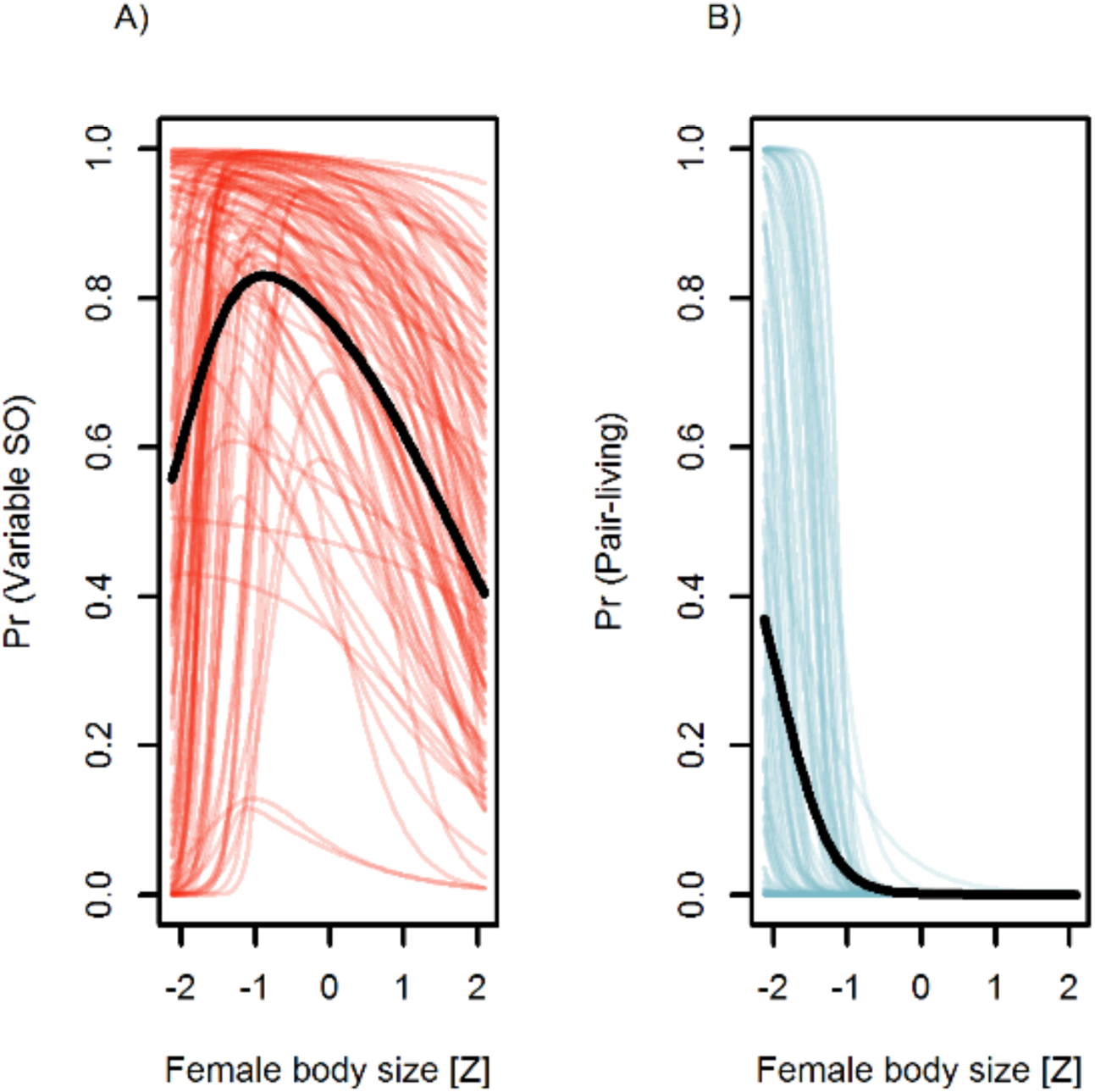
Probability of A) variable social organisation and B) pair-living as a function of female body size. Solid black lines indicate mean predicted values; thin lines represent 100 randomly drawn posterior samples to illustrate uncertainty.

The model for number of social organisations is summarized in Supplementary Materials S5. The predicted number of social organisations for the ancestor of all Artiodactyla was 1.73 (95% CI = 1.19 - 2.41), with no predictor influencing the number of social organisations at the 95% CI level. However, a decrease in number of social organisations with greater mean female body mass was relatively well supported (PP=0.91, consistent with Figure 2A). The prediction of fewer social organisations in open habitats compared to closed ones was again only weakly supported (PP=0.70). The phylogenetic signal was low (mean = 0.13, 95% CI = 1 e-5 – 0.29).

## 4. Discussion

Our dataset revealed that IVSO occurred in 75% of Artiodactyla species. For species showing IVSO, social organisation was variable within 62% of populations. These trends are consistent with previous descriptions of IVSO in artiodactyls [58] and other mammals including Carnivora [27% of species classified as ‘flexible’; 6], Eulipotyphla [43.8% of species with data; 21], Rodentia [16, 20], and strepsirrhine primates [60.5% of species with data; 22]. Mounting evidence of extensive IVSO in mammals challenges the common assumption in comparative studies that all species have only one social organisation [7, 59–61]. Failing to account for intraspecific variation will likely result in spurious conclusions about social evolution, slowing theoretical advancement [17, 25]. Using modern phylogenetic methods, we can now easily account for IVSO by analyzing data at the population rather than the species level. Moreover, greater effort should be made to build datasets from high-quality, primary sources rather than relying on secondary sources and taxonomic inference.

Our results change our understanding of social evolution. Both Jarman [18] and Pérez-Barbería et al. [13] assumed in their early comparative studies that the ancestral artiodactyl was socially monogamous (pair-living) with evolutionary transitions to polygyny and group-living. In contrast, Lukas & Clutton-Brock [7] argued that solitary living was the ancestral condition for most mammalian orders, including Artiodactyla. Contrary to these studies, our analysis estimated the ancestral social organisation to be variable, with possible transitions to both pair-living or group-living depending on body size, or sexual dimorphism and breeding seasonality, respectively (or possible unmeasured variables that cause variation in these factors). Thus, our study supports the argument that IVSO plays an important role in the evolution of mammalian social systems [6, 17].

Group-living and large body size are possible adaptations for artiodactyls living in open habitats [18, 40, 41] and to reduce predation risk [1]. Positive associations between group size and habitat openness have been observed in artiodactyls [62] and other mammals [e.g., 63]. Thus, we expected reduced IVSO in large bodied, group-living species in open habitats. In support of this hypothesis, both the probability of variable social organisation and the number of social organisations was low for species with large (and with very low) body mass but highest for species of intermediate body mass.

Contrary to expectations, IVSO did not increase with increasing number of habitat types, suggesting that IVSO is not the result of selection for habitat-specific social organisations. Furthermore, neither the probability of variable social organisation or group-living nor the number of social organisations differed between open (savannas and grasslands) and closed habitats (forests). Ecological conditions, such as the spatiotemporal distribution of food resources as a result of unpredictable and/or variable precipitation and temperature, may have a greater effect than habitat type on the social organisation, as was suggested for artiodactyls [64].

Reproductive competition changes seasonally in species that breed seasonally which in turn, can lead to greater variation in social organisation [36, 65]. Contrary to this expectation, artiodactyls that breed seasonally did not exhibit greater IVSO than non-seasonal breeders. Seasonal variation in local ecological conditions, such as the spatiotemporal variation in rainfall and food [17] may be more important predictors of IVSO than seasonality of breeding alone. The likelihood of IVSO increased with degree of sexual dimorphism. In polygynous systems, a large percentage of males may not breed [66]. Some males may live with other males in bachelor groups, increasing the prevalence of IVSO. In some species, such as African buffalo (*Syncerus caffer*) there is a rotation system in which breeding males join herds of breeding females for a period of time [67]. During this time, the males breed and fight, but then re-join bachelor groups to recover from the energetic costs of breeding [67].

In conclusion, our study demonstrated three major points regarding social evolution: 1) ancestral artiodactyl social organisation was variable and not pair-living, as was long assumed, 2) in artiodactyls, the frequency of IVSO increased with increasing sexual dimorphism and decreased with body size, and 3) taking IVSO into account and using a high-quality dataset significantly changes our understanding of social evolution. Our study should motivate future efforts to understand the importance of IVSO in animal social evolution.

## Data accessibility

R code and dataset available at https://github.com/adrianjaeggi/artiodactyl.socialorg.

## Authors’ contributions

L.D.H. and C.S. conceived of the project and contributed to manuscript writing. M.I.M. collected data and contributed to manuscript writing. A.J. conducted the statistical analysis and contributed to manuscript writing. M.F.B. provided insight into artiodactyls and contributed to manuscript writing.

## Acknowledgements

H. Klug and T. Gaudin provided helpful comments on the manuscript.

